# BIOFORMERS: A SCALABLE FRAMEWORK FOR EXPLORING BIOSTATES USING TRANSFORMERS

**DOI:** 10.1101/2023.11.29.569320

**Authors:** Siham Amara-Belgadi, Orion Li, David Yu Zhang, Ashwin Gopinath

## Abstract

Generative pre-trained models, such as BERT and GPT, have demonstrated remarkable success in natural language processing and computer vision. Leveraging the combination of large-scale, diverse datasets, transformers, and unsupervised learning, these models have emerged as a promising method for understanding complex systems like language. Despite the apparent differences, human language and biological systems share numerous parallels. Biology, like language, is a dynamic, interconnected network where biomolecules interact to create living entities akin to words forming coherent narratives. Inspired by this analogy, we explored the potential of using transformer-based unsupervised model development for analyzing biological systems and proposed a framework that can ingest vast amounts of biological data to create a foundational model of biology using BERT or GPT. This framework focuses on the concept of a ‘biostate,’ defined as a high-dimensional vector encompassing various biological markers such as genomic, proteomic, transcriptomic, physiological, and phenotypical data. We applied this technique to a small dataset of single-cell transcriptomics to demonstrate its ability to capture meaningful biological insights into genes and cells, even without any pre-training. Furthermore, the model can be readily used for gene network inference and genetic perturbation prediction.

## 1 Introduction

Large-scale, transformer-based models are revolutionizing natural language processing (NLP) and emerging as a new paradigm in the pursuit of artificial general intelligence[1]. Transformer models utilize a ‘self-attention’ mechanism to efficiently process and relate different input data parts. They are trained on vast datasets in a largely unsupervised fashion, enabling them to discern subtle patterns and relationships within the data[2]. Their success in NLP has positioned them as foundational tools for various applications and also given a blueprint for understanding other complex systems, like life sciences, purely from a corpus of data. In biology, the ‘language’ of living organisms is composed of biomolecules like DNA, RNA, proteins, and small molecules. These can be seen as the ‘words’ that interact in complex networks, akin to sentences in human language, forming structures like cells and tissues, analogous to paragraphs and chapters in a book. This perspective raises a fascinating question: How can we leverage techniques developed for human languages to gain foundational insights into biological systems?

Historically, biological research has focused on uncovering specific rules governing relationships between molecules, where discrete, isolated observations and theories drove understanding. This approach often involves studying individual biomolecules in isolation (or in small groups), much like trying to understand language by examining individual words (or sentences) without their complete contextual narrative. Over the last few decades, as the field has embraced the complexity and scale of living systems, it has become increasingly apparent that studying isolated components offers a limited view. This shift in perspective aligns well with the application of transformer-based approaches in AI. Transformer models have shown superior performance when trained on multiple, interconnected tasks rather than specific, isolated ones [3, 4, 5]. This characteristic makes them particularly suitable for studying inherently complex and interconnected biological systems. Simply put, just as these AI models have excelled in understanding the nuances of human language by analyzing words in the context of sentences and paragraphs, they hold promise in deciphering the ‘language’ of biological systems where molecules, cells, and tissues interact in complex ways.

Transformer models have already been applied to certain key areas of biology, most notably for protein design. Prominent examples include DeepMind’s AlphaFold [6] and Meta’s ESMFold [7], among others[8]. AlphaFold has been groundbreaking in predicting protein structures with remarkable accuracy, revolutionizing the field by providing insights into the 3D shapes of proteins, which are crucial for understanding their function. Similarly, Meta’s ESMFold contributes to this area by offering efficient and accurate protein structure predictions, further enhancing our ability to design and understand proteins. These applications of transformer models in protein design are akin to creating a perfect word in a language – they represent precise and highly specialized achievements. Yet, this success represents only a piece of the puzzle in understanding biological systems. Designing an ideal protein molecule is just one aspect; these molecules do not function in isolation but interact within complex systems.

While the feasibility of transformer-based models in studying systems biology remains largely unexplored, we are beginning to see promising attempts, particularly in the single-cell domain. This is primarily due to the scale of data available in single-cell experiments compared to other biological experiments. Single-cell RNA sequencing, for example, provides a wealth of data by analyzing thousands of individual cells[9] rather than averaging data across many cells. Thus, these single-cell experiments can be thought of as thousands of parallel experiments on nearly identical copies and can serve as a proxy for organism-level population-wide experiments, but at a markedly reduced cost[10], especially for model development.

These initial explorations have revealed a specific key challenge: the inability to transfer the insights gained by these models across different applications due to their distinct architectures. To overcome this, a wave of research initiatives [11, 12, 13, 14, 15] is underway, aiming to develop a foundational model that first deciphers latent information from unlabeled scRNA-seq data and then adapts this knowledge to various tasks. In scBERT[11], introduced by Yang and team in 2022, represents genes as tokens and employs an advanced transformer mechanism [16] to analyze over 16,000 gene tokens per cell. Another recent development, scGPT[14], introduces a variant of masked language modeling similar to autoregressive generation in natural language processing, iteratively predicting masked genes based on the model’s confidence level. These models focus primarily on uncovering the gene relationships within individual cells. They didn’t address how they could go beyond single cells and didn’t have features baked in to allow multi-omic integration or potential transfer learning between species.

In this work, we have proposed a framework, which we refer to as BioFormers, inspired heavily by scGPT [14] and scBERT [11] to operate on the biostate of sample and phenotypical information of a sample. The biostate, defined by us as a high-dimensional vector that includes various biological markers such as genomic data, proteomic data, transcriptomic data, physiological data, and phenotypical data. Further, the sample itself could be cells, tissue, organism, or longitudinal data. The core framework is transformer-based and designed to jointly optimize the sample and molecular embeddings. The model relies solely on self-attention, and we have approached the problem both as an encoder-only as well as a decoder-only design, with the preprocessing, tokenization, and embedding schema preserved. In the long run, we expect foundational models to be optimized at these multiple sample resolutions, i.e., cells, tissues, organisms, and longitudinal data, from which a mixture of models and fine-tuned models for an all-encompassing representation of biological systems. We demonstrate that our model is capable of getting meaningful insights with relatively small datasets without any pre-training. In a zero-shot setting, our model can perform slightly better than state-of-the-art on revealing biologically significant cell clusters, masked gene modeling, genetics perturbation prediction, as well as gene-network inference. We expect that with even a moderate amount of pre-training on larger datasets as well as higher quality datasets, which will be the focus of our follow-up study, the model is likely to

## 2 Methods

While the framework we have developed is designed to be general, in this work, we have specifically focused on applying it for scRNA sequencing. The transcriptomic data from single-cell RNA sequencing (scRNA-seq) is represented as a cell-gene matrix, denoted as *X* and belonging to the set of real numbers,*X* ∈ ℝ^*N ×G*^. In this matrix, each entry *X*_*i*,*j*_ represents the abundance RNA expressed from a gene *j* (where *j* is an index ranging from 0 to *G*) in cell *i* (where *i* is an index ranging from 0 to *N*), is a positive real number. We term this dataset the “raw matrix” in subsequent sections. Unlike natural language texts, the sequenced data in scRNA-seq is continuous, posing challenges for tasks like tokenization and modeling. In the following sections, we will introduce techniques for data processing and learning objectives to facilitate effective learning using transformer models.

**Figure 1:**
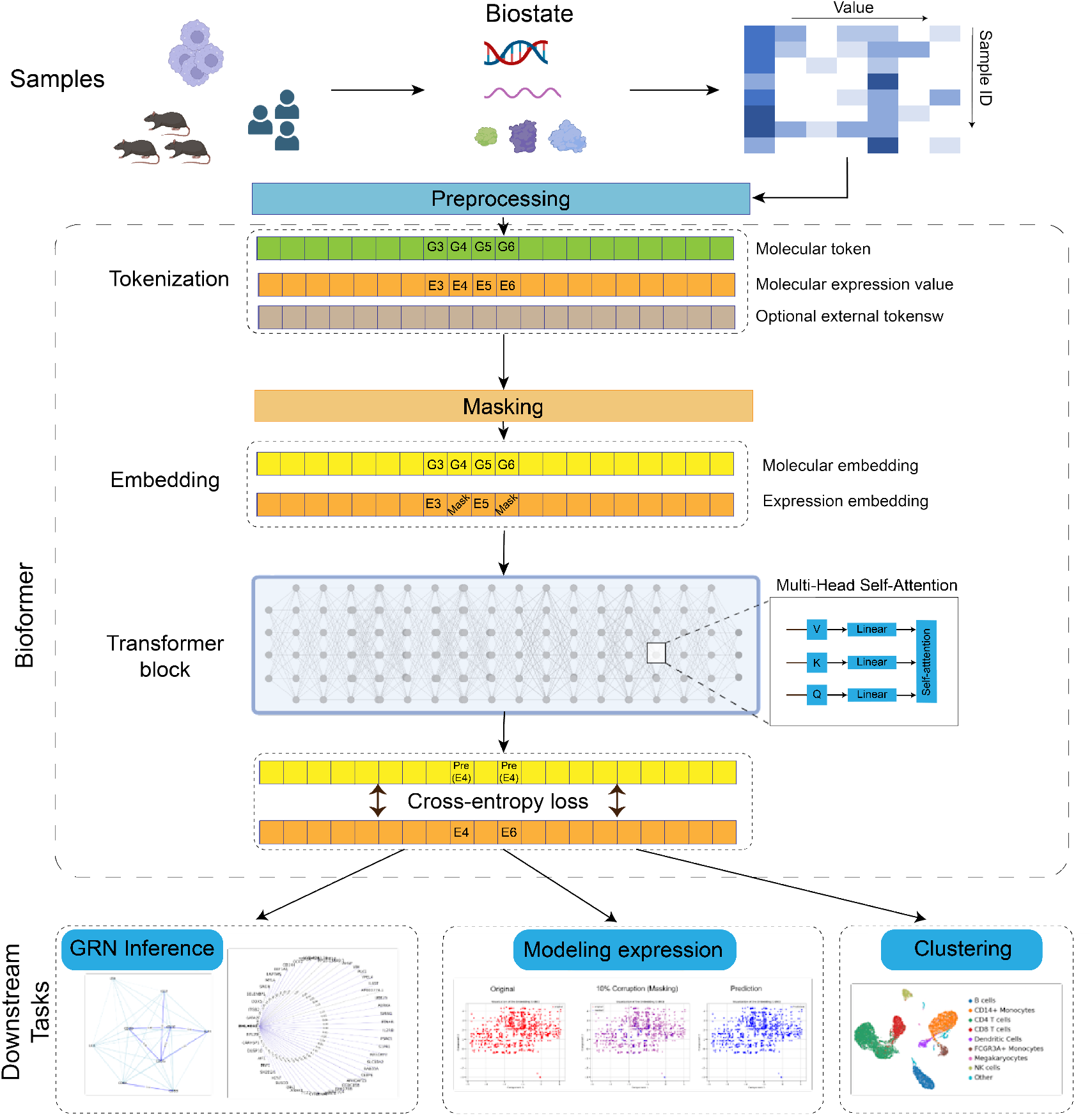
Model Schematic. BioFormers is a framework that is able to utilize various biological data, converting them through dedicated preprocessing techniques into a unified token representation. The tokens are then partially masked, converted into embeddings, and passed through multiple transformer blocks, before being decoded to predict the masked tokens from the unmasked ones. For downstream applications, BioFormers may retrieve general biological knowledge in a zero-shot learning process, and employ these knowledge in various biological tasks.

### 2.1 Input embeddings

The input to BioFormers consists of three components: (1) biomolecular tokens, (2) biomolecular expression values, and (3) optional external tokens. In this study, we have restricted to only using gene tokens and gene expression values as we use scRNA-seq data. The expression values are preprocessed from the raw matrix *X* with a slightly different procedure for each modeling task (see section X).

#### 2.1.1 Biomolecular Tokens

In our approach, we represent biomolecules as tokens, where each biomolecule, denoted as *bm*_*j*_, is assigned a unique integer *id*(*bm*_*j*_) from the complete token vocabulary. The input biomolecule tokens for each sample *i* is a fixed length vector *t*_*b*_*m*^(*i*)^ ∈ ℕ^*K*^ ,

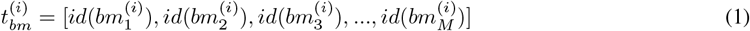

, where *M* is the present input length, usually set as the number of highly variable biomolecules used.

#### 2.1.2 Molecular Expression Values

The input of molecular expression values is converted into relative values from the raw counts *X*_*i*,*j*_. We adopted this approach because accurately modeling biomolecular expression poses significant challenges due to variations in absolute values across different assays [17]. These variations arise from differences in assay sensitivity and the low probability of detecting genes or molecules expressed at minimal levels. Data from various experiments can differ substantially in scale, even after undergoing standard preprocessing steps like normalization to a fixed sum and *log*1*p* transformation. Effectively, this means that identical absolute values can have different ‘semantic’ meanings in different experimental setups and batches. To address this issue, we propose two different binning **segment binning**, which involves ranking the expression count and then segmenting the rank, and **value binning**, which directly operates on the value.

##### Segment binning

For all nonzero expression counts of each sample (cell), we count the raw absolute values and make *B* several consecutive intervals [*b*_*k*_, *b*_*k*+1_], *k* ∈ 1, 2, …, *B*, where each interval range includes an equal 1*/B* portion of all expressed molecules. The computation itself is done sample-wise, and the interval edges *b*_*k*_ vary among samples. The converted expression value 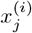 for each sample *i* is as follows,

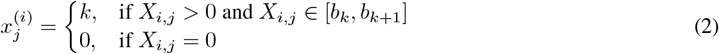

With segment binning, 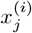will have more consistent semantic meaning across batches. For example, the value 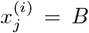 always implies that molecule *j* is one of the highest expressed molecules in the sample. The final input value vector for sample *i* is

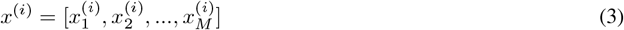

##### Value binning

We convert normalized and log-transformed expression counts into value bins. For raw expression count of gene *j* in cell *i, x*_*i*,*j*_, we first normalize across cells so that the sum of normalized expression counts of all cells equal the average sum of raw expression counts for each cell, resulting in normalized expression count:

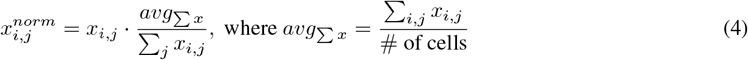

Then, we put the normalized expression counts into value bins by directly rounding at lower expression counts and doing log-transforming at higher expression levels. This provides the model with enough space to establish clear and detailed representation within the lower expression counts. Specifically, the binned expression value for gene *j* in cell *i*, 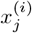, is as follows:

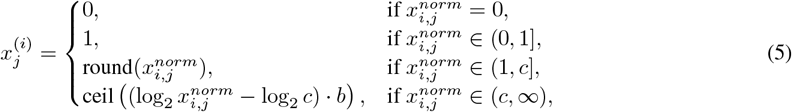

where *b* is the length of each log-transformed value bin, and *c* is the cutoff between lower and higher expression counts. With this binning, genes that fall within the same length-*b* segment after normalization and log transformation will have the same binned expression value 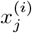, granting it consistency on the relative expression level across different genes within the same sequencing batch. During the experiments, we also train the model on value-binned data that are not normalized in order to explore the impact of normalization and the variance in the “semantic” meaning of gene expression counts.

#### 2.1.3 Optional External Tokens

Our framework allows for the inclusion of external tokens, which carry meta-information related to individual molecules. For instance, these tokens can indicate whether a molecule has undergone alterations in perturbation experiments. Additionally, they can carry a variety of molecular properties. For instance, if the token presents a particular protein, this might include the number of epitopes or details about available docking sites. In the scope of this study, we primarily utilized these tokens to denote molecular perturbations. However, the broader intent of integrating these tokens into our framework is to lay the groundwork for facilitating transfer learning in future research. We describe all external tokens as an input vector with the same dimension as the input molecules,

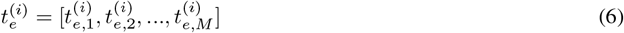

where 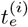 are integer indices represent external categories.

### 2.1.4 Embedding layers

We use standard embedding layers *emb*_*bm*_ for the biomolecular tokens and *emb*_*e*_ for the external tokens to map each token into an embedding vector of fixed-length *D*. Although the standard embedding layer is also applicable to the expression values since they are binned into a fixed set of *B* + 1 integers, we, by default, use fully connected layers, *emb*_*x*_. This has the benefit of easily modeling the consecutive nature of the value magnitudes. The final embedding *E*^(*i*)^ ∈ *R*^(*M×D*)^ of sample *i* is defined as

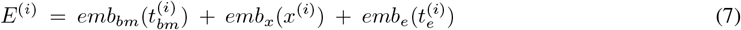

### 2.2 Molecular expression modeling

In this work, we have taken two approaches to biomolecular modeling. The first one is a transformer encoder model, as described by Vaswani et al.[2] and Devlin et al.[18], to encode the entire input embedding *h*^(*i*)^ as outlined in Equation 7. The transformer’s self-attention mechanism, which operates across the *M* embedding vectors in the input sequence, is particularly effective for learning interactions between molecules across various sample types. The output of the stacked transformer blocks is initially set as 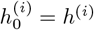, and then it is iteratively updated through the transformer blocks:

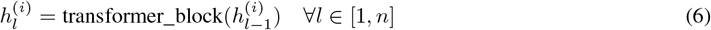

The final output, 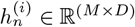, is used for both molecule-level and sample-level tasks. For molecule-level tasks, we directly apply task-specific heads to these learned embeddings. This specifically includes predictions of perturbed gene expression as well as Gene-Regulatory Network prediction. Regarding sample-level tasks, we first transform 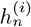into a consolidated sample embedding vector, described in the following subsection.

The second approach we pursued was inspired by masked-language modeling in natural language models (NLM). It employs masked molecule modeling to enhance the learning of cross-molecule relations. In each sample *i*, a proportion of molecules *j* and their processed expression values 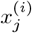 are randomly masked. The BioFormers model is then trained to accurately predict the molecular expression levels at these masked positions. This approach aids the model in effectively encoding co-expression patterns among molecule sets, ensuring that the attention mechanism of the model accurately represents the influence of each molecule *j* on another molecule *k*.

Formally, for the *n* binned molecule expression levels in sample *i*, denoted as 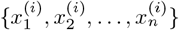, we mask *m × n* of these, where *m* is the masking ratio. Within these masked molecules, *p × m × n* have non-zero expression levels (i.e., 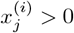), and (1 − *p*) ×*m* × *n* have zero expression levels. This distinction is crucial to counteract bias in scRNA-seq datasets, which typically feature a high ratio of zero expression levels, potentially leading the model towards a local minimum of outputting all zeros.

After the transformer processes the masked embedded input, its outputs are fed into a fully connected multi-layer perceptron (MLP). This MLP estimates the expression levels of the *m ×n* masked genes. We calculate these positions’ cross-entropy loss (CE) and employ an AdamW optimizer[19] for learning.

### 2.3 Sample representation

Our model treats each sample as a ‘sentence’ composed of molecules. To generate a sample representation 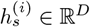, we integrate the learned molecule-level representation 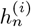. We further introduce a special token <cls> to represent the sample. This token is appended at the beginning of the input tokens sequence. During the transformer block processing, the model learns to pool information effectively, and the final embedding corresponding to the <cls> token position is used as the sample representation. This representation is typically the first row of 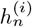, expressed as 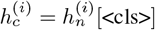, where [<cls>] denotes retrieving the row at the index of the <cls> token in the input.

## 3 Experiments and results

### 3.1 Modeling a single scRNA-seq experiment

In scRNA sequencing, one of the most crucial tasks is the classification of cells, with each cell viewed as an independent sample in our modeling framework. Cell clustering is the first step in identifying cell types and states. To assess the effectiveness of our model in capturing and preserving biological information, we initially focus on evaluating cell clustering. We do this by applying BioFormers to datasets with annotated cell types. This approach allows us to quantitatively measure how well the cell representations, as learned by the model, reflect and retain the underlying biological context. We tested one dataset re-processed by Gayoso et al. [20] composed of PBMC 8K (7,982 cells and 3,346 genes)

#### Setup

We utilized the SCANPY Python library [21] for preprocessing, which involved normalizing each cell by total gene counts, logarithmizing the data matrix using log1p, and selecting highly variable genes. To evaluate the cell embeddings, we employed biological conservation metrics as proposed by Luecken et al.[22], including Normalized Mutual Information (NMI), Adjusted Rand Index (ARI), and Average Silhouette Width (ASW). These metrics assess the alignment between the clusters derived from cell type data and the ground truth labels. For a comprehensive comparison, we introduced AvgBIO, the average of NMI, ARI, and ASW. Additionally, we benchmarked our BioFormers model against a baseline using highly variable gene (HVG) analysis, ensuring consistency across all methods by using the same set of HVGs. The output metrics were calculated using the implementation provided in Luecken et al. [22].

#### Result

The benchmarking results clearly illustrate that *BioFormers* performs well at the task of cell embedding extraction, outperforming existing models across all evaluated metrics. When assessed on the PBMC 8K dataset, the *BioFormers*’s **AvgBIO** score surpasses that of the Highly Variable Genes (HVG) baseline by a substantial margin of 5 to 19%. This improvement underscores the model’s capability to enhance biological signal detection and cell type differentiation, which is attributable to its sophisticated feature learning algorithms. While we have performed the analysis only with a single dataset, and much more analysis needs to be performed to validate these results, it is nonetheless very promising.

### 3.2 Gene expression modeling

The gene expression modeling experiment is a comprehensive test of our model’s ability to predict masked molecule gene levels, thereby assessing its proficiency in learning cross-molecule relationships and reconstructing any data within a cell’s context. We have adopted a suite of metrics beyond mere accuracy to evaluate the model’s performance holistically. For this purpose, we utilized a combined dataset from the PBMC 4k and 8k studies by Zheng et al.[23] and subsequently re-processed by Lopez et al.[24], with an applied filter for highly variable genes, resulting in a dataset comprising 11,990 cells and 2,000 HVGs. Post-filtering, we implemented a value binning technique with a bin size of *b* = 3 and a cutoff for high expression levels set at *c* = 8. The data was then divided, allocating 90% for training and 10% for validation.

#### Baseline

In establishing a comparative baseline, we employed Multi-variable Linear Regression. For each gene *j*, we trained a Linear Regression model using the *k* molecule *j*_1_, *j*_2_, …, *j*_*k*_ that exhibited the highest Pearson correlation with molecule *j* according to their value-binned expression levels in the training set. Through an exploration of various ensemble Linear Regression models with differing *k* values, our experiments indicated that sensitivity metrics begin to plateau when *k* reaches 300, whereas specificity metrics continue to decline. Consequently, we opted for *k* = 300 for our baseline Linear Regression model.

#### Setup

For value binning, we tested different setups of masking ratio *m* and nonzero ratio in the mask *p*, as well as whether or not the binning uses normalized raw expression counts. For the model, we use an 8-layer transformer encoder model with 8 self-attention heads per layer and a hidden state dimension of 512, trained through an AdamW optimizer using a cosine-reducing learning rate 2 × 10^−4^ with 2000 warm up steps. The training runs for 10 epochs on the training set.

#### Results

detailed in Table 2 utilize a suite of new metrics we have created to gauge the predictive accuracy of gene expression levels. ***Sensitivity*** measures the proportion of genes with nonzero expression levels that are correctly assigned to the precise bin. ***HEL Sensitivity*** evaluates the accuracy of predictions for High Expression Level genes, requiring predictions to be no more than one bin off for genes with expression levels above a certain threshold *c*. ***NPV-50*** assesses the accuracy of predicting unexpressed genes, specifically within a dataset with an equal distribution of zero and nonzero gene expression masks. The results from our experiments, particularly when masking 10% of the gene expression levels with half being nonzero expressions, demonstrated a notable increase in performance metrics: sensitivity for all genes and HEL genes increased by 30%, HEL Sensitivity rose by 9%, and NPV saw a 24% boost. These outcomes were superior to the baseline and illustrate the our model’s proficiency in capturing intricate relationships between genes and their expression levels, beyond mere numerical regressions, even without any domain-specific knowledge.

**Table 1:** Cell embedding results (PBMC 8K)

| Model | AvgBIO | ARI | NMI | ASW<sub>cell</sub> |
| --- | --- | --- | --- | --- |
| HVG | 0.626 | 0.624 | 0.733 | 0.522 |
| BioFormers | 0.819 | 0.786 | 0.861 | 0.810 |

**Table 2:**
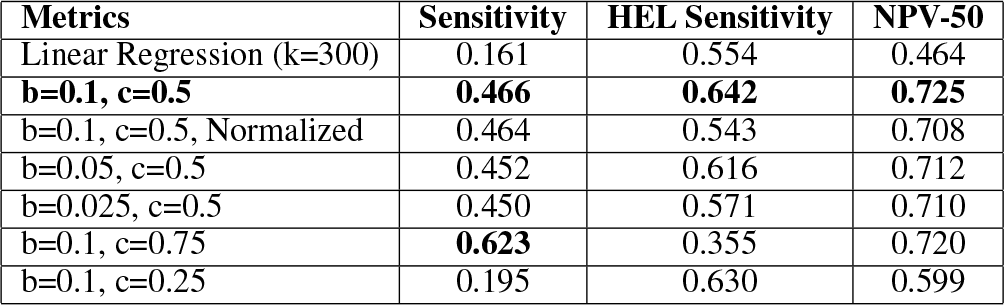
BioFormers models, while yielding slightly varying performances, all performs much better than Linear Regression on all metrics.

**Table 3:**
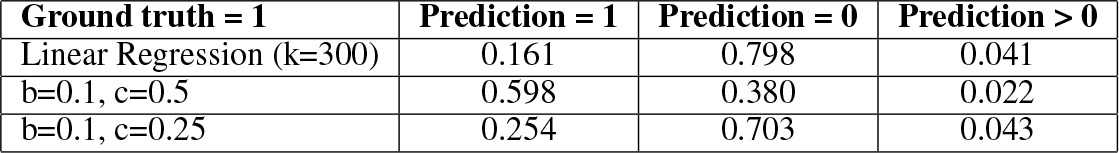
Heavily skewed data severely degrades Linear Regression Model performance on molecules with an actual expression of 1, but BioFormers Models are able to distinguish between zero and nonzero expressions successfully.

The Linear Regression Model’s suboptimal NPV performance can be partly attributed to the inherent bias in the dataset, where 89.2% of the raw molecular expression counts are 0, and another 7.5% are 1. This skew in the data likely biases the training, leading to a predisposition in the model towards predicting a zero expression level, particularly when the actual expression is 1, thereby diminishing its sensitivity. Conversely, the optimal BioFormers model showcases enhanced sensitivity for genes with an actual expression of 1. Moreover, the model with *b* = 0.1 and *c* = 0.75 demonstrates high sensitivity, likely due to an over-representation of nonzero molecules in the mask and a preponderance of 1s among the nonzero genes, resulting in a bias towards predicting 1s for actual expressions of 0s and 1s, which is evident from the reduced specificity. These findings indicate that future training should focus on mitigating biases induced by the nature of scRNA-seq datasets, striving for a more balanced representation of molecular expression across zero, low, and high levels.

To gauge the impact of database quality and size on our model’s performance, we retrained it using the meticulously curated Perturb-seq dataset by Adamson et al. [25]. The outcomes, delineated in Table 4, reveal a substantial enhancement in model efficacy, with an 18% rise in HEL Sensitivity. However, we did observe an overall dip in sensitivity which we ascribe to a misalignment between the dataset’s normalization technique and our value binning methodology, mirroring the performance reduction seen in Table 2, attributable to analogous normalization disparities.

**Table 4:** The same model trained on a larger dataset yields notable improvement, particularly in HEL Sensitivity.

| Metrics | Training Set Size | Sensitivity | HEL Sensitivity |
| --- | --- | --- | --- |
| PBMC 4k + 8k | 10997 | 0.466 | 0.642 |
| Adamson Perturb-seq | 69560 | 0.430 | **0.823** |

**Table 5:** Perturbation generation results.

| Model | (DE) $\text{corr}$ | (DE) $\text{corr}(\Delta)$ | ALL $\text{corr}$ | ALL $\text{corr}(\Delta)$ |
| --- | --- | --- | --- | --- |
| MLP | 0.94 | 0.72 | 0.991 | 0.656 |
| BioFormers | 0.96 | 0.740 | 0.991 | 0.632 |

### 3.3 Post perturbation expression prediction

Combining gene-editing techniques and single-cell RNA sequencing has enabled high-throughput experiments that uncover cellular reactions to various genetic alterations. These advancements hold great promise for uncovering new gene interactions and advancing regenerative medicine. Nonetheless, the vast combinatorial space of potential gene perturbations swiftly surpasses the number of experiments that can practically be conducted, thus constraining its utility. Machine learning, however, offers a solution by learning from the cellular responses of known perturbations and predicting outcomes of untested ones. Such datasets are uniquely suited to test the performance of BioFormers, whose self-attention mechanism captures the relationships between perturbed genes and the subsequent expression changes in other genes. We evaluated BioFormers in a scenario designed to predict gene expression following perturbations. For this task, our benchmark dataset was the Perturb-seq dataset by Adamson et al. [25], which was preprocessed by Roohani et al. [26]. The Perturb-seq dataset encompasses 87 single-gene perturbations, with approximately 100 cells per perturbation and a control set of at least 7,000 unperturbed cells.

#### Setup

In the one gene perturbation experiments, we adhered to the preprocessing protocol established by Roohani et al. [26], which includes normalization by total molecular counts, log transformation of the data, selection of 5,000 highly variable molecules, and inclusion of any perturbed molecules otherwise omitted. To ensure the validity of our perturbation prediction task, we meticulously separated the perturbations such that none of the test perturbations were exposed to the model during training, guaranteeing no overlap between training and test cells in terms of perturbations. The accuracy of perturbation prediction was measured by the Pearson correlation (corr) between the predicted and actual molecular expression values post-perturbation. In addition, we utilized a variation of the Pearson metric that assesses the change in expression post-perturbation relative to the control (denoted as *corr(*∆)). Our evaluation extended to various molecular sets, specifically the complete molecular set (ALL) and the top 20 deferentially expressed molecules (DE), resulting in four distinct evaluation metrics: *corr* and *corr(*∆) for both (ALL) and (DE) molecular sets.

#### Results

In benchmarking against the multi-layer perceptron (MLP) baseline, our BioFormers model consistently registers the highest correlation with the true perturbed molecular expression values across all evaluated metrics. Given that approximately 50% of molecular expression counts remain unchanged post-perturbation, attributable to either low sensitivity of the assay to those molecules rates or inherently low molecular expression the metrics focusing on differentially expressed molecules (DE) offer a more compelling assessment. Notably, BioFormers exhibits enhancements in the correlation metric (∆) for the top differentially expressed molecules, a measure that is arguably of the greatest significance.

### 3.4 GRN inference

Gene Regulatory Networks (GRNs) are pivotal in mediating biological processes, with transcription factors, co-factors, enhancers, and target genes interacting in complex ways. Traditional methods for inferring GRNs often utilize static gene expression correlations or pseudo-time estimates to approximate causal relationships, as outlined by Aditya Pratapa et al. [27]. However, attention scores derived from our models offer a dynamic approach to understanding GRNs. These scores pinpoint key genes influencing specific outcomes, with higher scores indicating greater importance.

**Figure 2:**
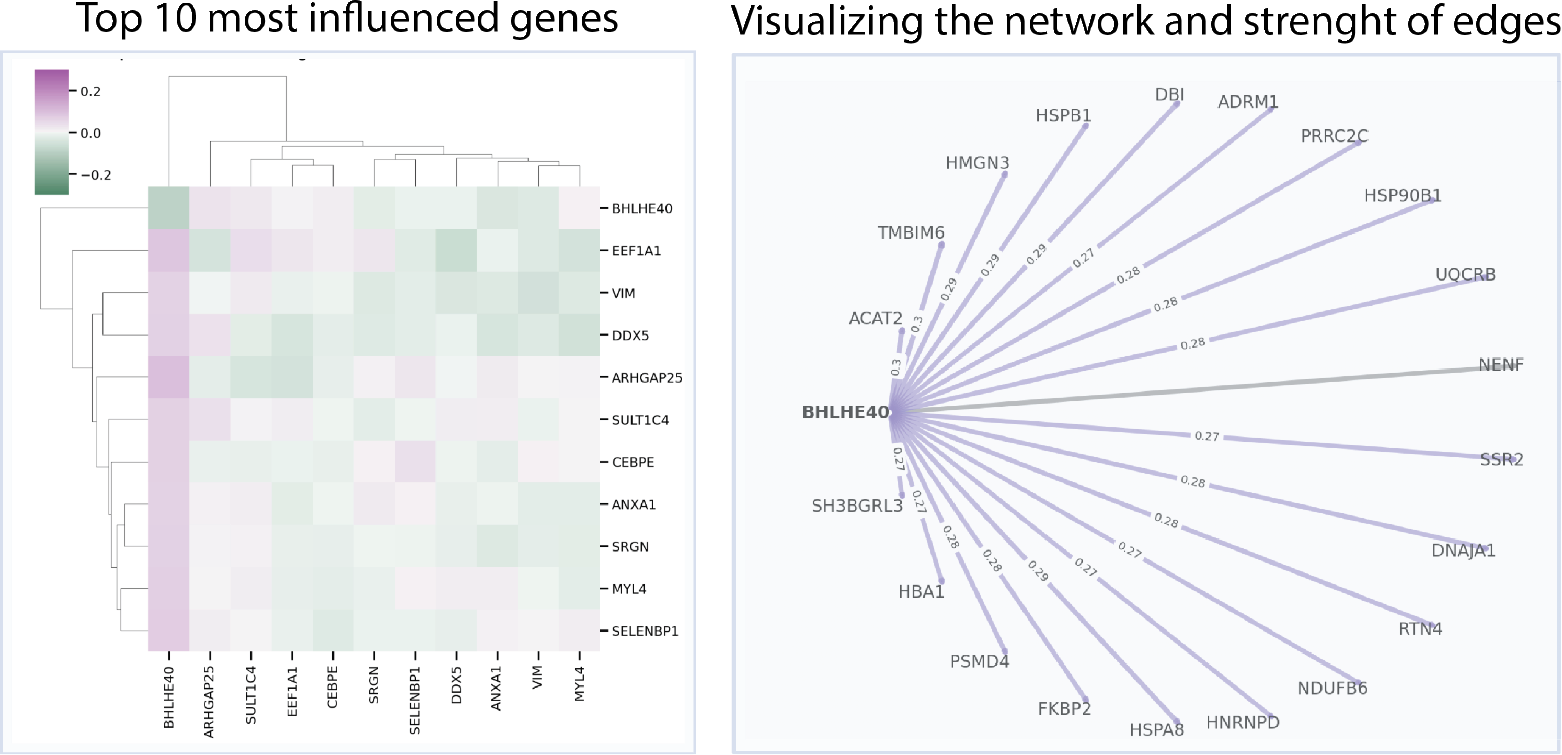
Heatmap attention score for the 10 most important Genes for BHLHE40 transcription factor repression, and gene target network graph of the top 20 genes validated in the CHIP-Atlas database

Additionally, they reveal gene-to-gene correlations, with mutual high attention scores suggesting strong interactions, which can be condition-specific when aggregated across different conditions. Furthermore, genes can be clustered by the similarity of their attention scores, reflecting commonalities in function or expression. In a zero-shot framework, we derive GRNs from model-generated gene embeddings based on cosine similarities and identify gene programs from clusters containing five or more genes. This method allows the visualization of gene connectivity within specific programs, such as those including CD3 and CD8 clusters, where strong connections, denoted by high cosine similarities and illustrated in blue, are observable among genes like CD3D, CD3E, CD3G, CD8A, and CD8B.

While our approach is adept at discerning gene connectivities within clusters, it does exhibit variability in attention scores across different training iterations. For instance, disparate results were observed when training the transformer encoder model multiple times; one instance failed to grasp the connectivity within the CD3 cluster. Another iteration accurately identified interactions within the Histone cluster, especially between the linker gene H1-2 and the H2B cluster gene H2BC12 [28]. However, the same iteration missed capturing connections that another iteration did successfully, like those between HBZ and HBA1 within the Human Alpha Globin cluster. These inconsistencies underscore the need for refined training methodologies to ensure comprehensive learning of gene connectivities and address the current shortcomings.

To identify key regulators or drivers of gene expression, we utilize attention scores computed through a self-attention mechanism for each gene within the dataset. The genes with the highest attention scores are flagged as pivotal in predicting the expression levels of others. Subsequent pathway analysis or gene ontology examination of these key genes sheds light on the biological pathways or processes they predominantly influence. Further, by consulting a database of known transcription factor-gene interactions, we pinpoint the transcription factors regulating these influential genes. Utilizing the Adamson perturbation dataset, we contrast the impacted genes between transcription factor repression conditions (perturbed) and control. This method of attention-based GRN inference transcends perturbation studies, extending to broader comparisons such as healthy versus diseased or undifferentiated versus differentiated cell states. We aggregate attention scores per condition, summing across all input data batches to obtain average attention scores for each condition. This sets the stage for selecting the most influenced genes and examining network changes post-perturbation. Validation is conducted against CHIP-Atlas, which includes targets of the transcription factor BHLHE40, allowing us to assess the concurrence between top selected genes and known targets. Finally, we visualize the network, elucidating the strength of the connections among genes.

## 4 Conclusions

Inspired by the prowess of Large Language Models (LLMs) in uncovering complex patterns in natural language through predominantly unsupervised methods [1], we ventured to assess the efficacy of similar approaches for deciphering the ‘language’ of biology. We introduce a transformer-based framework, termed BioFormer, which leverages a ‘self-attention’ mechanism to interpret high-dimensional biological data. We demonstrated the merit of our approach by applying it to learn the underlying patterns of several single-cell RNA-seq datasets. We consistently achieved superior results compared to traditional methods in tasks such as cell clustering and masked gene prediction. Our cell clustering performance surpassed state-of-the-art methods by 30.8%, and our masked gene prediction improved upon the linear regression baseline by 15.8% to 189.4%, depending on the chosen metric. We also successfully applied our technique to elucidate gene regulatory networks solely from the data in a zero-knowledge context, without the need for additional system knowledge or pre-training.

Looking ahead, we aim to expand our pre-training to encompass larger, more diverse datasets, including multi-omic data, diseased states, perturbations, and temporal sequences [29]. BioFormer’s framework includes joint molecular and sample tokenization, enabling generative pre-training on heterogeneous data, alongside optional external tokens for facilitating transfer learning across sample types. Our next steps involve pre-training BioFormer on comprehensive bulk RNA-seq data from rats, followed by fine-tuning on longitudinal time-series data. We anticipate that our approach will contribute to the expanding corpus of research that employs transformer-based models to comprehend biological data, laying the groundwork for a foundational model of biology.

## Code availability

The code for producing the baseline results is publicly available on our <u>GitHub repository</u>

## Author contributions

AG and DYZ conceptualized the study and directed the research. SA, OL, AG designed an performed the experiments as well as analyzed the data. SA, OL, and AG wrote the paper with input from all authors.

## Notes

https://github.com/BiostateAI/

### Competing Interest Statement

All authors are employees of Biostate AI. Additionally, DYZ declares competing interest in the form of consulting for and significant equity ownership in Pupil Bio, NuProbe Global, and Torus Biosystems. AG declares competing interest in the form of consulting for and significant equity interest in Iuno and Argome. All other authors have no other competing interests.

## References

[1] OpenAI. GPT-4 Technical Report. 2023. Publisher: arXiv Version Number: 3.

[2] Ashish Vaswani, Noam Shazeer, Niki Parmar, Jakob Uszkoreit, Llion Jones, Aidan N. Gomez, Lukasz Kaiser, and Illia Polosukhin. Attention Is All You Need. 2017. Publisher: arXiv Version Number: 7.

[3] Suchin Gururangan, Ana Marasović, Swabha Swayamdipta, Kyle Lo, Iz Beltagy, Doug Downey, and Noah A. Smith. Don’t Stop Pretraining: Adapt Language Models to Domains and Tasks. 2020. Publisher: arXiv Version Number: 3.

[4] Chi Sun, Xipeng Qiu, Yige Xu, and Xuanjing Huang. How to Fine-Tune BERT for Text Classification? In Maosong Sun, Xuanjing Huang, Heng Ji, Zhiyuan Liu, and Yang Liu, editors, Chinese Computational Linguistics, volume 11856, pages 194–206. Springer International Publishing, Cham, 2019. Series Title: Lecture Notes in Computer Science.

[5] XiPeng Qiu, TianXiang Sun, YiGe Xu, YunFan Shao, Ning Dai, and XuanJing Huang. Pre-trained models for natural language processing: A survey. Science China Technological Sciences, 63(10):1872–1897, October 2020.

[6] John Jumper, Richard Evans, Alexander Pritzel, Tim Green, Michael Figurnov, Olaf Ronneberger, Kathryn Tunyasuvunakool, Russ Bates, Augustin Žídek, Anna Potapenko, Alex Bridgland, Clemens Meyer, Simon A. A. Kohl, Andrew J. Ballard, Andrew Cowie, Bernardino Romera-Paredes, Stanislav Nikolov, Rishub Jain, Jonas Adler, Trevor Back, Stig Petersen, David Reiman, Ellen Clancy, Michal Zielinski, Martin Steinegger, Michalina Pacholska, Tamas Berghammer, Sebastian Bodenstein, David Silver, Oriol Vinyals, Andrew W. Senior, Koray Kavukcuoglu, Pushmeet Kohli, and Demis Hassabis. Highly accurate protein structure prediction with AlphaFold. Nature, 596(7873):583–589, August 2021.

[7] Zeming Lin, Halil Akin, Roshan Rao, Brian Hie, Zhongkai Zhu, Wenting Lu, Nikita Smetanin, Robert Verkuil, Ori Kabeli, Yaniv Shmueli, Allan Dos Santos Costa, Maryam Fazel-Zarandi, Tom Sercu, Salvatore Candido, and Alexander Rives. Evolutionary-scale prediction of atomic level protein structure with a language model. preprint, Synthetic Biology, July 2022.

[8] John B. Ingraham, Max Baranov, Zak Costello, Karl W. Barber, Wujie Wang, Ahmed Ismail, Vincent Frappier, Dana M. Lord, Christopher Ng-Thow-Hing, Erik R. Van Vlack, Shan Tie, Vincent Xue, Sarah C. Cowles, Alan Leung, João V. Rodrigues, Claudio L. Morales-Perez, Alex M. Ayoub, Robin Green, Katherine Puentes, Frank Oplinger, Nishant V. Panwar, Fritz Obermeyer, Adam R. Root, Andrew L. Beam, Frank J. Poelwijk, and Gevorg Grigoryan. Illuminating protein space with a programmable generative model. Nature, November 2023.

[9] Dragomirka Jovic, Xue Liang, Hua Zeng, Lin Lin, Fengping Xu, and Yonglun Luo. Single-cell RNA sequencing technologies and applications: A brief overview. Clinical and Translational Medicine, 12(3):e694, March 2022.

[10] John M. Ashton, Hubert Rehrauer, Jason Myers, Jacqueline Myers, Michelle Zanche, Malene Balys, Jonathan Foox, Chistopher E. Mason, Robert Steen, Marcy Kuentzel, Catharine Aquino, Natàlia Garcia-Reyero, and Sridar V. Chittur. Comparative Analysis of Single-Cell RNA Sequencing Platforms andMethods. Journal of Biomolecular Techniques : JBT, 32(4):3fc1f5fe.3eccea01, December 2021.

[11] Fan Yang, Wenchuan Wang, Fang Wang, Yuan Fang, Duyu Tang, Junzhou Huang, Hui Lu, and Jianhua Yao. scBERT as a large-scale pretrained deep language model for cell type annotation of single-cell RNA-seq data. Nature Machine Intelligence, 4(10):852–866, September 2022.

[12] Jing Gong, Minsheng Hao, Xin Zeng, Chiming Liu, Jianzhu Ma, Xingyi Cheng, Taifeng Wang, Xuegong Zhang, and Le Song. xTrimoGene: An Efficient and Scalable Representation Learner for Single-Cell RNA-Seq Data. preprint, Bioinformatics, March 2023.

[13] Hongru Shen, Xilin Shen, Jiani Hu, Jilei Liu, Chao Zhang, Dan Wu, Mengyao Feng, Meng Yang, Yang Li, Yichen Yang, Wei Wang, Qiang Zhang, Jilong Yang, Kexin Chen, and Xiangchun Li. Generative pretraining from large-scale transcriptomes: Implications for single-cell deciphering and clinical translation. preprint, Bioinformatics, February 2022.

[14] Haotian Cui, Chloe Wang, Hassaan Maan, Kuan Pang, Fengning Luo, and Bo Wang. scGPT: Towards Building a Foundation Model for Single-Cell Multi-omics Using Generative AI. preprint, Bioinformatics, May 2023.

[15] Christina V. Theodoris, Ling Xiao, Anant Chopra, Mark D. Chaffin, Zeina R. Al Sayed, Matthew C. Hill, Helene Mantineo, Elizabeth M. Brydon, Zexian Zeng, X. Shirley Liu, and Patrick T. Ellinor. Transfer learning enables predictions in network biology. Nature, 618(7965):616–624, June 2023.

[16] Krzysztof Choromanski, Valerii Likhosherstov, David Dohan, Xingyou Song, Andreea Gane, Tamas Sarlos, Peter Hawkins, Jared Davis, Afroz Mohiuddin, Lukasz Kaiser, David Belanger, Lucy Colwell, and Adrian Weller. Rethinking Attention with Performers. 2020. Publisher: arXiv Version Number: 4.

[17] Abhishek Sarkar and Matthew Stephens. Separating measurement and expression models clarifies confusion in single-cell RNA sequencing analysis. Nature Genetics, 53(6):770–777, June 2021.

[18] Jacob Devlin, Ming-Wei Chang, Kenton Lee, and Kristina Toutanova. BERT: Pre-training of Deep Bidirectional Transformers for Language Understanding. 2018. Publisher: arXiv Version Number: 2.

[19] Ilya Loshchilov and Frank Hutter. Decoupled Weight Decay Regularization. 2017. Publisher: arXiv Version Number: 3.

[20] Adam Gayoso, Romain Lopez, Galen Xing, Pierre Boyeau, Valeh Valiollah Pour Amiri, Justin Hong, Katherine Wu, Michael Jayasuriya, Edouard Mehlman, Maxime Langevin, Yining Liu, Jules Samaran, Gabriel Misrachi, Achille Nazaret, Oscar Clivio, Chenling Xu, Tal Ashuach, Mariano Gabitto, Mohammad Lotfollahi, Valentine Svensson, Eduardo Da Veiga Beltrame, Vitalii Kleshchevnikov, Carlos Talavera-López, Lior Pachter, Fabian J. Theis, Aaron Streets, Michael I. Jordan, Jeffrey Regier, and Nir Yosef. A Python library for probabilistic analysis of single-cell omics data. Nature Biotechnology, 40(2):163–166, February 2022.

[21] F. Alexander Wolf, Philipp Angerer, and Fabian J. Theis. SCANPY: large-scale single-cell gene expression data analysis. Genome Biology, 19(1):15, December 2018.

[22] Malte D. Luecken, M. Büttner, K. Chaichoompu, A. Danese, M. Interlandi, M. F. Mueller, D. C. Strobl, L. Zappia, M. Dugas, M. Colomé-Tatché, and Fabian J. Theis. Benchmarking atlas-level data integration in single-cell genomics. Nature Methods, 19(1):41–50, January 2022.

[23] Grace X. Y. Zheng, Jessica M. Terry, Phillip Belgrader, Paul Ryvkin, Zachary W. Bent, Ryan Wilson, Solongo B. Ziraldo, Tobias D. Wheeler, Geoff P. McDermott, Junjie Zhu, Mark T. Gregory, Joe Shuga, Luz Montesclaros, Jason G. Underwood, Donald A. Masquelier, Stefanie Y. Nishimura, Michael Schnall-Levin, Paul W. Wyatt, Christopher M. Hindson, Rajiv Bharadwaj, Alexander Wong, Kevin D. Ness, Lan W. Beppu, H. Joachim Deeg, Christopher McFarland, Keith R. Loeb, William J. Valente, Nolan G. Ericson, Emily A. Stevens, Jerald P. Radich, Tarjei S. Mikkelsen, Benjamin J. Hindson, and Jason H. Bielas. Massively parallel digital transcriptional profiling of single cells. Nature Communications, 8(1):14049, January 2017.

[24] Romain Lopez, Jeffrey Regier, Michael B. Cole, Michael I. Jordan, and Nir Yosef. Deep generative modeling for single-cell transcriptomics. Nature Methods, 15(12):1053–1058, December 2018.

[25] Britt Adamson, Thomas M. Norman, Marco Jost, Min Y. Cho, James K. Nuñez, Yuwen Chen, Jacqueline E. Villalta, Luke A. Gilbert, Max A. Horlbeck, Marco Y. Hein, Ryan A. Pak, Andrew N. Gray, Carol A. Gross, Atray Dixit, Oren Parnas, Aviv Regev, and Jonathan S. Weissman. A Multiplexed Single-Cell CRISPR Screening Platform Enables Systematic Dissection of the Unfolded Protein Response. Cell, 167(7):1867–1882.e21, December 2016.

[26] Yusuf Roohani, Kexin Huang, and Jure Leskovec. GEARS: Predicting transcriptional outcomes of novel multi-gene perturbations. preprint, Bioinformatics, July 2022.

[27] Aditya Pratapa, Amogh P. Jalihal, Jeffrey N. Law, Aditya Bharadwaj, and T. M. Murali. Benchmarking algorithms for gene regulatory network inference from single-cell transcriptomic data. Nature Methods, 17(2):147–154, February 2020.

[28] William F. Marzluff, Preetam Gongidi, Keith R. Woods, Jianping Jin, and Lois J. Maltais. The Human and Mouse Replication-Dependent Histone Genes. Genomics, 80(5):487–498, November 2002.

[29] Wei Chen, Yi Chai, Qi Jiang, Eva Y. Miao, Ashwin Gopinath, and David Yu Zhang. High Frequency Longitudinal RNAseq Reveals Temporally Varying Genes and Recovery Trajectories in Rats. preprint, Molecular Biology, November 2023.

